# Inhibiting monocyte migration reduces arterial thrombosis and facilitates thrombolysis

**DOI:** 10.1101/2024.07.21.604499

**Authors:** Hee Jeong Jang, Jiwon Kim, Ha Kim, Taesu Kim, Jinyong Chung, Sebastian Cremer, Marvin Krohn-Grimberghe, Dawid Schellingerhout, Matthias Nahrendorf, Dong-Eog Kim

## Abstract

**Background:** Monocytes contribute to the initiation and propagation of venous thrombosis. Little is known about the roles monocytes play in arterial thrombosis, the cause of stroke and MI.

**Methods:** We investigated how chemokine receptor 2 (CCR2) knockout (^-/-^) affects platelet function, blood coagulation, thrombus volume, and thrombolytic susceptibility in 627 mice with FeCl_3_-mediated carotid arterial thrombosis: three CX3CR1-GFP mice, 326 C57BL/6 mice, and 288 CCR2^-/-^ mice. We performed i) intravital microscopy imaging of leukocyte recruitment to carotid thrombus, ii) flow cytometry-based quantification of leukocytes in blood vs. thrombus and leukocyte-platelet aggregates in blood, iii) platelet aggregometry, iv) coagulation assays, v) micro-computed tomography (microCT)-based thrombus imaging using gold nanoparticles after tissue plasminogen activator (tPA) therapy with or without either a) CCR2 siRNA pretreatment (for 3 days) or post-treatment or b) clopidogrel pretreatment (for 7 days), and vi) histology including scanning electron microscopy.

**Results:** Intravital microscopy and flow cytometry showed that both neutrophils and monocytes were recruited to the acute arterial thrombus, as observed 30 minutes post-thrombosis. Platelet function tests demonstrated platelet aggregation to be lower in the whole blood of CCR2^-/-^ mice (vs. C57BL/6 mice) but not in their leukocyte-free platelet rich plasma, suggesting this platelet dysfunction is cell-mediated. Flow cytometry experiments revealed lower numbers of monocyte – platelet aggregates (MPAs), a marker of platelet activity, in the blood of CCR2^-/-^ mice, compared to C57BL/6 mice. Blood levels of FXIII and monocyte levels of FXIII-A were increased after carotid thrombosis in C57BL/6 mice but not CCR2^-/-^ mice. Further, *in vivo* microCT and histology, respectively, showed that CCR2^-/-^ mice had smaller and more porous thrombi with less fibrin cross-linking, compared with C57BL/6 mice. MicroCT also demonstrated that tPA-mediated thrombolysis was faster in CCR2^-/-^ mice and CCR2 siRNA-treated mice, compared to C57BL/6 mice. In addition, clopidogrel had a greater effect on inducing thrombus formation with smaller sizes after FeCl3 application, while CCR2^-/-^ had a greater effect on dissolving thrombus faster after tPA administration.

**Conclusions:** CCR2 antagonism decreases platelet aggregation and reduces FXIII levels in blood and monocytes, thus driving arterial thrombosis towards the generation of a relatively small, porous, more lysable clot.

**Novelty and Significance:** *What Is Known?:*

- In venous thrombosis, monocytes cooperate with platelets and neutrophils to promote thrombus formation; however, few studies have investigated the roles of monocytes in arterial thrombosis.
- An *in vitro* study using arterial blood samples from patients with acute coronary syndrome showed that P-selectin on activated platelets upregulated monocyte chemokine receptor 2 (CCR2) expression, which was higher in monocyte-platelet aggregates (MPAs) than in monocytes without platelets.
- In a mouse study using an *in vivo* model of FeCl_3_-induced arterial thrombosis, the time to thrombotic occlusion was prolonged in CC chemokine ligand 2 (CCL2) knockout (^-/-^) mice compared with wild-type mice.

*What New Information Does This Article Contribute?:*

- This study is the first to demonstrate that CCR2^-/-^-mediated monocyte deficiency exerts antithrombotic and tissue plasminogen activator (tPA)-facilitating effects in acute arterial thrombosis.
- CCR2^-/-^ mice had a) decreased platelet aggregability, probably due in part to reduced post-thrombosis MPA numbers, and b) lower blood and monocyte levels of Factor XIII (FXIII), compared to those of C57BL/6 mice.
- Micro-computed tomography (microCT) thrombus imaging showed that i) CCR2^-/-^ mice, compared to C57BL/6 mice, had smaller thrombi, with more porosity and less fibrin cross-linking on histology and ii) CCR2^-/-^ or CCR2 siRNA facilitates tPA-mediated thrombolysis. The majority of ischemic strokes occur due to thromboembolic occlusion of cerebral arteries. Tissue plasminogen activator (tPA), which is the only FDA-approved drug for acute ischemic stroke, has only a moderate efficacy rate regarding early recanalization, which is closely associated with thrombus volume and thrombolytic resistance. Monocytes, in cooperation with platelets and neutrophils, contribute to thrombus formation in venous thrombosis. To date, only a few studies have detailed potential prothrombotic roles for monocytes in arterial thrombosis. In the present study, using a mouse model of FeCl_3_-mediated carotid artery thrombosis and high-resolution microCT thrombus imaging, we demonstrate that CCR2^-/-^-mediated monocyte deficiency exerts antithrombotic and tPA-facilitating effects. Compared to C57BL/6 mice, CCR2^-/-^ mice exhibited decreased platelet aggregability along with lower numbers of circulating MPAs (a marker of platelet activity) and reduced monocyte recruitment to arterial thrombi. Moreover, CCR2^-/-^ mice had lower levels of both circulating FXIII and monocyte FXIII-A, which are known to strengthen and stabilize thrombi. In line with these findings, i) CCR2^-/-^ mice had smaller, more porous thrombi with less fibrin cross-linking and ii) tPA-mediated thrombolysis was faster in CCR2^-/-^ mice or with CCR2 siRNA.

## Introduction

Stroke is a leading cause of death and long-term disability. About 80% of strokes are ischemic strokes, which occur when blood flow to the brain is obstructed due to thromboembolic occlusion of cerebral arteries. Despite the proven efficacy of tissue plasminogen activator (tPA)-mediated thrombolysis, up to ∼60% of patients die or become dependent after the intravenous recanalization therapy,^1^ due in part to only a moderate rate of early recanalization efficacy: 30%-40% overall, with <∼10% toward the most proximal occlusions.^2–4^ tPA efficacy is closely associated with thrombus burden^5^ and thrombolytic resistance.^6^

Research on venous thrombosis has shown that monocytes cooperate with platelets and neutrophils, contributing to thrombus formation and propagation.^7^ However, only a few studies have investigated the roles of monocytes in arterial thrombosis.^8^ A study using patient arterial blood samples revealed that platelets and monocytes collaborate to induce thromboinflammation during acute coronary syndrome.^9^ This *in vitro* study showed that P-selectin on activated platelets upregulated monocyte chemokine receptor 2 (CCR2) expression, which was higher in monocyte-platelet aggregates (MPAs) than in monocytes without platelets.^9^ In work using a mouse model of FeCl_3_-induced arterial thrombosis, platelets expressed CC chemokine ligand 2 (CCL2, also known as monocyte chemoattractant protein-1), which increased platelet aggregation, activation, and α-granule secretion.^10^ The time to thrombotic occlusion was prolonged in CCL2 knockout (^-/-^) mice compared with wild-type mice,^10^ and the authors suggested that further research on arterial thrombosis using CCR2^-/-^ mice is required.^10^ Moreover, the current literature contains limited information on FeCl_3_-mediated arterial thrombosis-related changes in platelet function and coagulation assays in both C57BL/6 and CCR2^-/-^ mice.

Our present study seeks to bridge these knowledge gaps. We utilized a mouse model of FeCl_3_-mediated carotid artery thrombosis to investigate CCR2 knockout effects on a) platelet function as determined by coagulation assays and b) arterial thrombus volume and tPA-mediated thrombolysis rate *in vivo* as measured using micro-computed tomography (microCT)-based high-resolution direct thrombus imaging.^11,12^

## Methods

### Animals and *in situ* carotid artery thrombosis model

This study was approved by the Institutional Animal Care and Use Committee at Dongguk University Ilsan Hospital and conducted in accordance with the NIH Research Rigor and Reproducibility guidelines. Twelve-week-old C57BL/6 mice (n = 339), CCR2 -/- mice (n = 290), CX3CR1-GFP mice (n = 3) were purchased from DBL Co. (Incheon, South Korea) and the Jackson Laboratory (Bar Harbor, ME, USA) and fed *ad libitum* in a specific pathogen-free and climate-controlled environment maintained at 20°C and 40-50% humidity, with 12 hours of light per 24-hour period. Mice were anesthetized with 2% isoflurane delivered through an inhalation mask. Experienced researchers (J. Kim and H.J. Jang) formed *in situ* carotid thrombi, as previously published,^12,13^ by exposing the vessel via a midline neck incision, dissecting the perivascular tissue, and then applying a strip of 1 × 1 mm^2^ filter paper (grade 42; Whatman, Oxon, UK) soaked in 4 μL 10% FeCl_3_ to the left common carotid artery (CCA) for 10 minutes.

### Experimental Groups

After exclusion of five mice due to experimental failures, a total of 622 animals were included in the final analyses. In the first set of experiments, 56 mice underwent intravital microscopy imaging of leukocyte recruitment to arterial thrombi (n=3 CX3CR1-GFP knock-in mice), flow cytometry analysis of leukocyte populations in blood and thrombus (n=19 C57BL/6 mice and 19 CCR2^-/-^ mice), and RNA expression analysis of thrombus-containing carotid arteries vs. non-thrombosed contralateral vessels (n=15 C57BL/6 mice). Subsequently, only C57BL/6 mice and CCR2^-/-^ mice were used, as follows: 73 and 74 mice for the second set of experiments using platelet aggregometry to assess platelet function (n=5-20/group); 35 and 26 mice for the third set of experiments on visual inspection-based (12 and 12; n=6/group) and flow cytometry-based (23 and 14; n=7-13/group) counting of MPAs and neutrophil-platelet aggregates (NPAs); 115 and 120 mice for the fourth set of experiments determining blood / monocyte coagulation parameters (n=4-21/group) and tail bleeding test (n=7-9/group); 9 and 9 mice for the fifth set of histology experiments including Carstair’s staining (n=3/group), Masson’s trichrome (MT) staining (n=3/group), and scanning electron microscopy (SEM) imaging (n=3/group); and 70 and 40 mice for the sixth set of experiments using the *in situ* carotid thrombosis model and high-resolution microCT thrombus imaging (n=6-27/group). Animals were randomly assigned to experimental groups by either H.J. Jang or S.-K. Lee. Outcome assessors, data analysts, and operators were all kept blinded to allocation.

### Intravital two-photon microscopy of leukocyte recruitment to thrombus

Intravital microscopy (IVIM-C; IVIM, Seoul, South Korea) was performed using CX3CR1-GFP mice to observe the accumulation leukocytes, including green fluorescent monocytes, in arterial thrombi. Two hours after intravenous injection of SeTau-647-labeled Ly6G antibody (1 mg/kg in 100 μL saline; IVIM, Seoul, South Korea) to label neutrophils, baseline imaging was performed, followed by the induction of carotid thrombosis. Imaging was then conducted immediately after the procedure and again at 30 minutes.

### Flow cytometry analysis of leukocyte populations in blood and thrombus

To assess leukocyte populations in blood by multicolor flow cytometry (BD^®^ LSR II Flow Cytometer; BD biosciences, San Jose, CA, USA), whole blood (200∼300μl) was collected from retro-orbital plexus to a 5ml EDTA blood tube (Soyagreentec, Seoul, South Korea). Thrombi were extracted from the carotid artery, and cell suspensions were prepared for multicolor flow cytometry to measure leukocyte populations. Seeking the mechanism of leukocyte accumulation in arterial thrombus (i.e., active recruitment vs. passive entrapment), we compared leukocyte contents in thrombus vs. blood from each mouse at 30 min and 4 days after carotid thrombosis. The following purified or conjugated antibodies (BioLegend, San Diego, CA, USA) were used: purified anti-FcγRIII/II (CD32/16, clone 2.4G2); FITC-labeled anti-Ly6C (Gr-1); PE-labeled anti-CD90.2 (Thy-1.2); PerCP-labeled anti-CX3CR1 (SA011F11); PE-Cy7-labeled anti-F4-80 (BM8); Brilliant Violet (BV) 421-labeled anti-CD115 (AF598); BV 605-labeled anti-CD19 (HIB19); BV 711-labeled anti-CD45 (30-F11); APC-labeled anti-Ly6G (1A8); AF700-labeled anti-NK1.1 (PK136); and APC-Cy7-labeled anti-CD11b (M1/70). Dead cells were gated out by AmCyan labeling. If monocyte recruitment to carotid thrombi was observed, RNA was isolated from the thrombus-containing arteries to identify factors potentially associated with this recruitment.^14^ We assessed the expression of chemokines, cytokines, and vascular adhesion molecules in 15 C57BL/6 mice (n=4-5/biomarker).^14^ The contralateral healthy vessel served as control tissue. mRNA was extracted using the RNeasy Mini Kit (Qiagen, Hilden, Germany) according to the manufacturer’s protocol. Then, mRNA was transcribed to cDNA using the High Capacity RNA-to-cDNA Kit (Applied Biosystems, Foster City, CA, USA). TaqMan primers (Applied Biosystems, USA) were utilized for quantitative analysis, and results were expressed as Cycle threshold values normalized to the housekeeping gene GAPDH (Glyceraldehyde 3-phosphate dehydrogenase).

### Platelet aggregometry

Whole blood (∼500 μL) was collected from the retro-orbital plexus either 30 minutes post-thrombosis or without inducing thrombosis, to account for potential effects of the thrombosis and related procedure, as previously reported.^15^ Platelet-rich plasma (PRP) was prepared using a double-spin method as previously described, as follows: whole blood was added to a centrifuge tube containing 3.2% (w/v) trisodium citrate (9:1 v/v mixture). The tubes were centrifuged at 800 rpm for 15 min, resulting in three layers: a platelet-poor plasma (PPP) layer at the top of the tube, a PRP layer in the middle, and an erythrocyte layer at the bottom. The supernatant yellow plasma (PRP layer) was centrifuged at 3,500 rpm for 15 min to concentrate the platelets. Then, the platelet-rich pellet was resuspended in 0.5∼1 ml of PPP, and the suspension was collected as PRP. Finally, the platelet count was adjusted to 5×10^7^ platelets/250µl with PPP.

Platelet aggregability was measured for using anticoagulated whole blood in a citrate tube and using PRP in a 1.5 ml microcentrifuge tube (SARSTEDT, Nümbrecht, Germany) by using a Lumi-aggregometer with Aggro/link 8 software (Chrono-Log Corporation, Havertown, PA, USA), as we previously reported.^15^ Platelet aggregation was induced by adding thrombin (1 unit), collagen (2 μg/mL), adenosine diphosphate (ADP; 10 μmol/L), or arachidonic acid (AA, 0.5 mmol/L) to 250 μL of sample (1:1 prediluted with saline). Immediately after adding one of these agonists, the following aggregometry data were obtained for the first 6 minutes, as we previously published:^15^ the area under the aggregation curve (AUC; Ohm × minute, a measure of overall platelet aggregation), amplitude (A6; amplitude at 6 minutes in Ohm, a measure of the extent of platelet aggregation), and the slope of the aggregation curve (a measure of how quickly platelets aggregate).^16^

### Quantification of leukocyte-platelet aggregates

Immunofluorescence staining of leukocytes and platelets was performed for whole blood (200 μL in a citrate tube) obtained either 30 minutes post-thrombosis or without inducing thrombosis, with or without adding platelet agonists (arachidonic acid [AA] 0.5 mmol/L, collagen 2 μg/mL, or adenosine diphosphate [ADP] 10 μmol/L) by using rat anti-Mouse CD16/CD32 Fc block (BD biosciences, Heidelberg, Germany) and the following purified or conjugated antibodies (BioLegend, San Diego, CA, USA): PE-labeled anti-CD62P (RMP-1) antibody and 488-labeled anti-CD115 (AFS98) antibody. After fixation and red blood cell lysis with BD 5× Lyse/Fix Buffer (BD Biosciences, San Jose, CA, USA), samples were centrifuged at 3,500 rpm for 15 min, and the pellets were resuspended in Cell Staining Buffer (BioLegend, San Diego, CA, USA) for fluorescence microscopy. For flow cytometry, purified Rat Anti-Mouse CD16/CD32Fc block (BD Biosciences, Heidelberg, Germany) and the following purified or conjugated antibodies (BioLegend, San Diego, CA, USA) were used: APC-labeled anti-CD41 (MWReg30); PE/Cy7-labeled anti-CD62P (RMP-1); BV 421-labeled anti-CD115 (AFS98); PerCP/Cy5.5-labeled anti-CD11b (M1/70); PE-labeled anti-Ly6G (1AB); FITC-labeled anti-Ly6C (HK1.4). After fixation and red blood cell lysis with BD 5× Lyse/Fix Buffer (BD Biosciences, San Jose, CA, USA), samples were centrifuged at 3,500 rpm for 15 min, and the pellets were resuspended in Cell Staining Buffer (BioLegend, San Diego, CA, USA). Neutrophils and monocytes were identified as CD11b^+^ Ly6G^+^ (pan-myeloid) cells and CD11b^+^ CD115^+^ Ly6C^+^ cells, respectively. Pan-platelets and activated platelets were identified as CD41^+^ cells and CD62P^+^ (P-selectin^+^) cells, respectively. NPAs were identified as CD11b^+^ Ly6G^+^ CD41^+^ CD62P^+^ cells. MPAs were identified as CD11b^+^ CD115^+^ Ly6C^+^ CD41^+^ CD62P^+^ cells. BD LSRFortessa™ (BD Biosciences, San Jose, CA, USA) and FlowJo version 10.8.1 (TreeStar, Ashland, OR USA) were used for flow cytometry and data analysis, respectively.

### Tail bleeding test

As previously reported,^15,17^ a distal 1 cm segment of the tail was amputated under anesthesia and monitored in 37℃ saline for 20 minutes, even if bleeding stopped, in order to detect any rebleeding.

### Coagulation assays

Blood coagulation assays were performed using whole blood (∼400 μL) collected through either retro-orbital or cardiac puncture in mice, either 30 minutes post-thrombosis or without thrombosis, as previously reported.^15^ Blood was mixed with 3.2% sodium citrate buffer (Medicago, Quebec City, Canada) at a 9:1 ratio. After plasma was isolated, prothrombin time (PT) and activated partial thromboplastin time (aPTT) were measured using a coagulation analyzer (Copresta 2000, Sekisui Medical, Tokyo, Japan). For PT, thromboplastin and calcium were added to the plasma, and clotting time was measured. For aPTT, phospholipid, ellagic acid, and calcium chloride were added to the plasma, and clotting time was measured.^18^ Enzyme-linked immunosorbent assay (ELISA) kits were used to measure plasma concentrations of factor III (FIII, tissue factor), FVII, FIX, FX, FXII (LSBio, Seattle, WA, USA), FXIII (Abbexa, Cambridge, UK), and thrombin-antithrombin complex (TAT; Abcam, Cambridge, UK). In addition, procoagulant activities of FXa, FXIIIa, and thrombin were measured using activity assay kits (BioVision, Mountain View, CA, USA).

To measure FXIII-A and tissue factor in monocytes, a microbead kit (CD115 Microbead Kit; Miltenyi Biotec, Bergisch Gladbach, Germany) was used to isolate these cells from the whole blood of C57BL/6 and CCR2^-/-^ mice, either 30 minutes post-thrombosis or without thrombosis. Total protein was extracted from the isolated monocytes by adding 100 μL of radioimmunoprecipitation buffer (Thermo Fisher Scientific, Waltham, MA, USA) supplemented with protease and phosphatase inhibitors (Thermo Fisher Scientific, Waltham, MA, USA). The mixture was briefly vortexed and then incubated on ice for 30 minutes. After incubation, the samples were centrifuged at 12,000 rpm for 15 minutes at 4°C to pellet cellular debris. The supernatant containing the extracted protein was carefully collected and stored at -80°C. Then, monocyte levels of FXIII-A and tissue factor were measured using ELISA (LSBio, Seattle, WA, USA) and western blotting, respectively. For western blotting, we used primary antibodies against β-actin (Cell Signaling Technology, Danvers, MA, USA) and tissue factor (Antibodies-online, Limerick, PA, USA), along with a protein detection and analysis instrument (ProteinSimple; Bio-techne Korea, Gwacheon, South Korea).^19,20^

### Histology

Carotid arteries containing thrombus were retrieved 30 minutes after thrombosis induction, fixed with 4% formaldehyde for 24 hours, and embedded in paraffin. Carstair’s staining (StatLab, McKinney, TX, USA) and Masson’s trichrome (MT) staining (ScyTek, Logan, UT, USA) were performed on 4 μm-thick serial sections obtained using a microtome (RM2235, LEICA, Wetzlar, Germany). Representative microscopy (Olympus, Tokyo, Japan) images were captured using the Cellsens standard imaging software (Olympus, Tokyo, Japan).^7,21,22^

### SEM

Carotid arteries containing thrombi were fixed in a glutaraldehyde buffer (2.5% in 0.1M phosphate buffer, pH 7.4; Ladd Research, PA, USA), followed by dehydration, critical point drying, resin infiltration, and embedding processes. Next, specimens were mounted on aluminum stubs and sputter coated with a layer of platinum, using a sputter coater (208HR; Cressington Scientific Instruments, Watford, UK). SEM was then performed using a field emission scanning electron microscope (Hitachi S-4800 FE-SEM; Hitachi, Tokyo, Japan) at 3 kV.

### *In vivo* direct thrombus imaging-based quantification of thrombus volume to assess antithrombotic effects of CCR2 knockout, CCR2 siRNA, tPA, and antiplatelet therapy

After fibrin-targeted glycol-chitosan-coated gold nanoparticles (fib-GC-AuNPs) were synthesized, microCT-based (NFR Polaris-G90, NanoFocusRay, Jeonju, South Korea) thrombus imaging was performed ∼30 minutes after carotid thrombosis and ∼5 minutes after intravenous injection of 200 μL fib-GC-AuNPs (2 mg/kg) via the penile vein in 70 C57BL/6 mice and 40 CCR2^-/-^ mice, as previously reported^11,12,23^ with some modifications: 500 ms per frame, 65 kVp, 60 uA, 360 views, high magnification, 512 × 512 reconstruction matrix, and 600 slices. Additional imaging was performed for 43 of the 70 C57BL/6 mice and 18 of the 40 CCR2^-/-^ mice that received no additional treatment (n=9 C57BL/6 and 10 CCR2^-/-^ mice) or received either 3-day pretreatment (n=14) or a single intravenous post-treatment dose (n=12) of either CCR2-siRNA (n=13) or vehicle (n=13) (in C57BL/6 mice) or 7-day oral pretreatment with either clopidogrel (2 mg Plavix^®^ dissolved in 100 uL PBS for 5 C57BL/6 mice and 5 CCR2^-/-^ mice; Sanofi, Paris, France) or saline (for 3 mice of each strain). This additional imaging was serially performed at 1, 2, 3, and 24 hrs after intravenous tPA therapy (20mg/kg Actilyse^®^, 10% bolus + 90% infusion over 30 minutes; Boehringer Ingelheim Korea, Seoul, South Korea) that was initiated immediately after baseline imaging at ∼30 minutes. The last pretreatment dose was administered immediately before thrombosis induction. Thrombus volume was quantified using Image J (NIH, Bethesda, MD, USA), as previously reported,^11,12^ by an experienced researcher (J. K.) blinded to the experimental groups.

### Statistical analyses

Data are presented as the mean ± standard deviation (SD). Comparisons of means between two groups were performed using the Mann-Whitney U test. Comparisons involving more than two groups were conducted using the one-way ANOVA, followed by Tukey’s *post hoc* tests for multiple comparisons. To compare repeated measures among time-points or groups, we used linear mixed models with random intercepts and pairwise *post hoc* tests with Sidak’s adjustment for multiple comparisons. A P value of < 0.05 was considered statistically significant. Statistical analyses were performed using SPSS 18.0 (SPSS Inc., Chicago, IL, USA) and Prism 9.5.1 (GraphPad Software, San Diego, CA, USA).

## Results

### CCR2^-/-^ mice showed less monocyte recruitment to carotid artery thrombus, compared to C57BL/6 mice

Intravital microscopy of CX3CR1-GFP mice showed real-time accumulation of monocytes and neutrophils in acute carotid thrombus generated by FeCl3 application (Fig. 1A). Flow cytometry at 30 minutes and 4 days after thrombosis demonstrated recruitment of monocytes and neutrophils to carotid thrombus (Fig. 1B-C and Supplementary Fig. 1). As expected, circulating monocytes were more abundant in C57BL/6 mice, compared to CCR2^-/-^ mice, at both 30 minutes and 4 days. Likewise, there were more monocytes in thrombi of C57BL/6 mice, compared to CCR2^-/-^ mice, at both 30 minutes and 4 days. Circulating neutrophils were reduced in C57BL/6 mice compared to CCR2^-/-^ mice at 30 minutes, while neutrophil counts in thrombi were non-significantly lower in C57BL/6 mice. At 4 days, however, C57BL/6 mice had more circulating neutrophils compared to CCR2^-/-^ mice, whereas neutrophil counts in thrombi were lower in C57BL/6 mice. Lymphocyte counts in thrombus and blood showed no significant inter-group differences at either 30 minutes or 4 days, except that circulating lymphocyte counts were significantly higher in C57BL/6 mice, compared to CCR2^-/-^ mice, at 30 minutes (Fig. 1B-C and Supplementary Fig. 1). Moreover, at 30 minutes (Fig. 1C), C57BL/6 mice had notably (∼20-fold) higher thrombus-blood ratios for both neutrophil counts (4.2±0.9) and monocyte counts (2.9±1.0) compared with the ratios of lymphocyte counts (0.2±0.1). Likewise, in CCR2^-/-^ mice, the thrombus-blood leukocyte ratios were substantially higher for both neutrophils (2.7±0.6) and monocytes (5.2±5.4), compared with the ratio for lymphocytes (0.2±0.2). At 4 days (Fig. 1B), in both C57BL/6 and CCR2^-/-^ mice, total leukocyte counts in thrombi were ∼10-fold lower (compared to 30-minute leukocyte counts) while monocyte counts were increased. Thus, the thrombus-blood ratios at 4 days (Fig. 1C) were significantly higher for monocytes than for neutrophils, and again lowest for lymphocytes, in both strains. Taken together, these findings suggest active recruitment of neutrophils and monocytes to the site of acute arterial thrombosis;^24^ such recruitment may occur via cytokine (CCL2, CCL5, CCL7 CXCL2, and CX3CR1) and/or chemokine (IL-6 and IL-1b)-related mechanisms (Supplementary Fig. 2). Subsequent experiments were performed in the hyperacute (i.e., 30-minute) setting.

**Fig 1.**
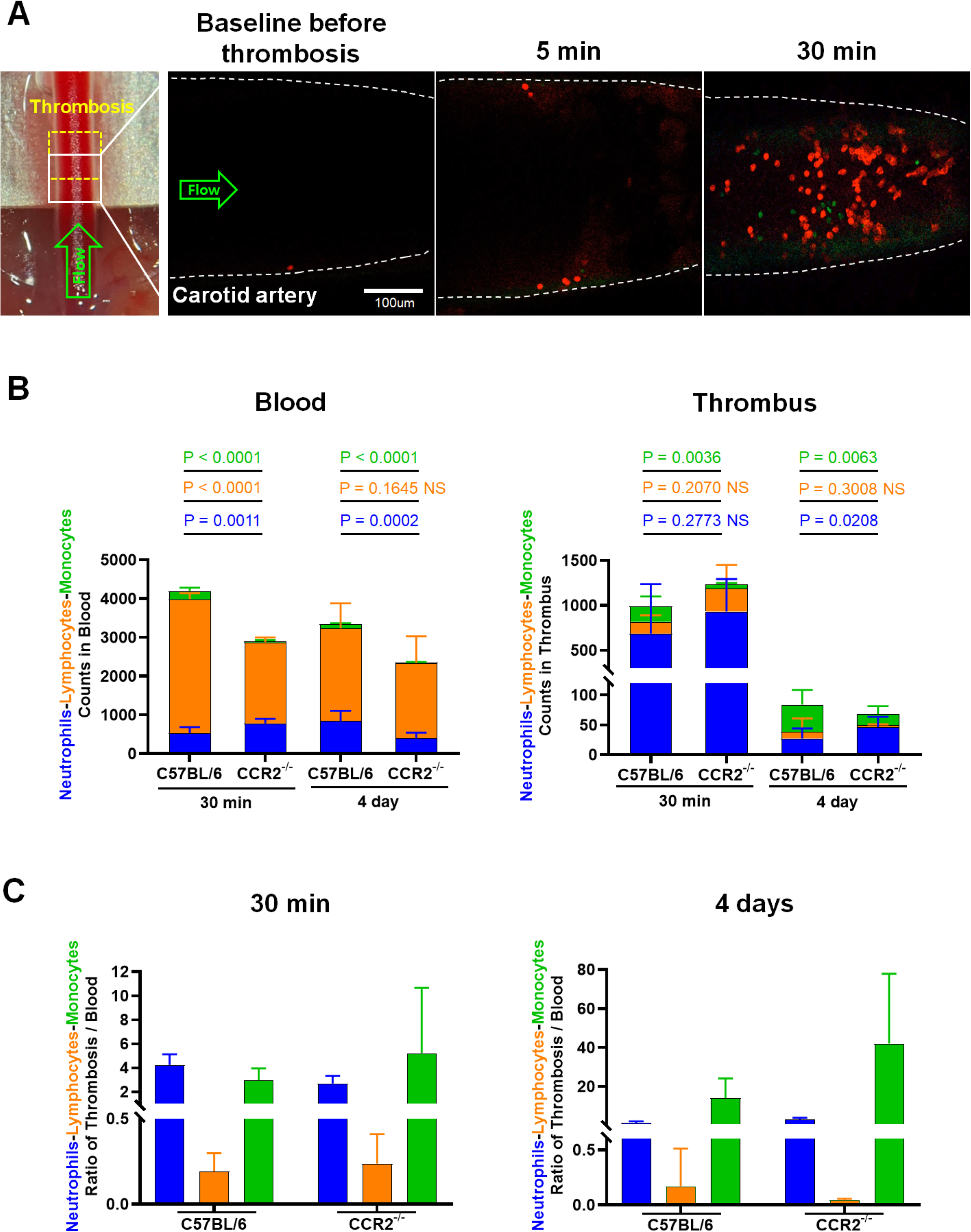
Monocytes and neutrophils accumulate in arterial thrombus. **A**. Intravital microscopy images of GFP^+^ monocytes (green) and SeTau-647-labeled Ly6G^+^ neutrophils (red) in the carotid thrombus of a representative CX3CR1-GFP knock-in mouse 30 minutes after FeCl3 application (yellow dotted square) on the carotid artery for 10 minutes. **B**. Flow cytometry analysis of leukocyte counts in blood and thrombus at 30 minutes (left) and 4 days (right) after carotid thrombosis in C57BL/6 and CCR2^-/-^ mice. **C**. Thrombus-to-blood ratios of leukocyte counts at 30 minutes (left) and 4 days (right), calculated using the flow cytometry data for **B**. Note the substantially higher thrombus-blood ratios for neutrophils and monocytes (vs. lymphocytes) in both C57BL/6 and CCR2-/- mice, suggesting active recruitment of circulating myeloid cells to thrombi. The most prevalent leukocytes in blood are lymphocytes, while in thrombi, neutrophils predominate. Monocytes are relatively abundant in thrombi (vs. blood) of C57Bl/6 mice (vs. CCR2-/- mice).

### In whole blood, but not leukocyte-deficient PRP, platelet aggregability was lower in CCR2^-/-^ mice compared to C57BL/6 mice

Platelet function tests of arterial whole blood samples (Fig. 2A) revealed the response to collagen stimulation had significantly lower values in terms of the AUC of the aggregation curve in non-thrombosed CCR2^-/-^ mice, compared to non-thrombosed C57BL/6 mice. In response to thrombin stimulation, the AUC were distinctly lower in non-thrombosed CCR2^-/-^ mice than in non-thrombosed C57BL/6 mice. There were no noteworthy inter-group differences in response to either AA or ADP stimulation. Further platelet function tests showed no significant difference in how arterial whole blood samples from the thrombosed CCR2^-/-^ and C57BL/6 mouse groups responded to either collagen or thrombin stimulation (Fig. 2B). However, the response to ADP stimulation showed lower AUC values in thrombosed CCR2^-/-^ mice, compared to thrombosed C57BL/6 mice. In addition, the response to AA stimulation tended to show lower AUC and slope values in thrombosed CCR2^-/-^ mice than in thrombosed C57BL/6 mice.

**Fig 2.**
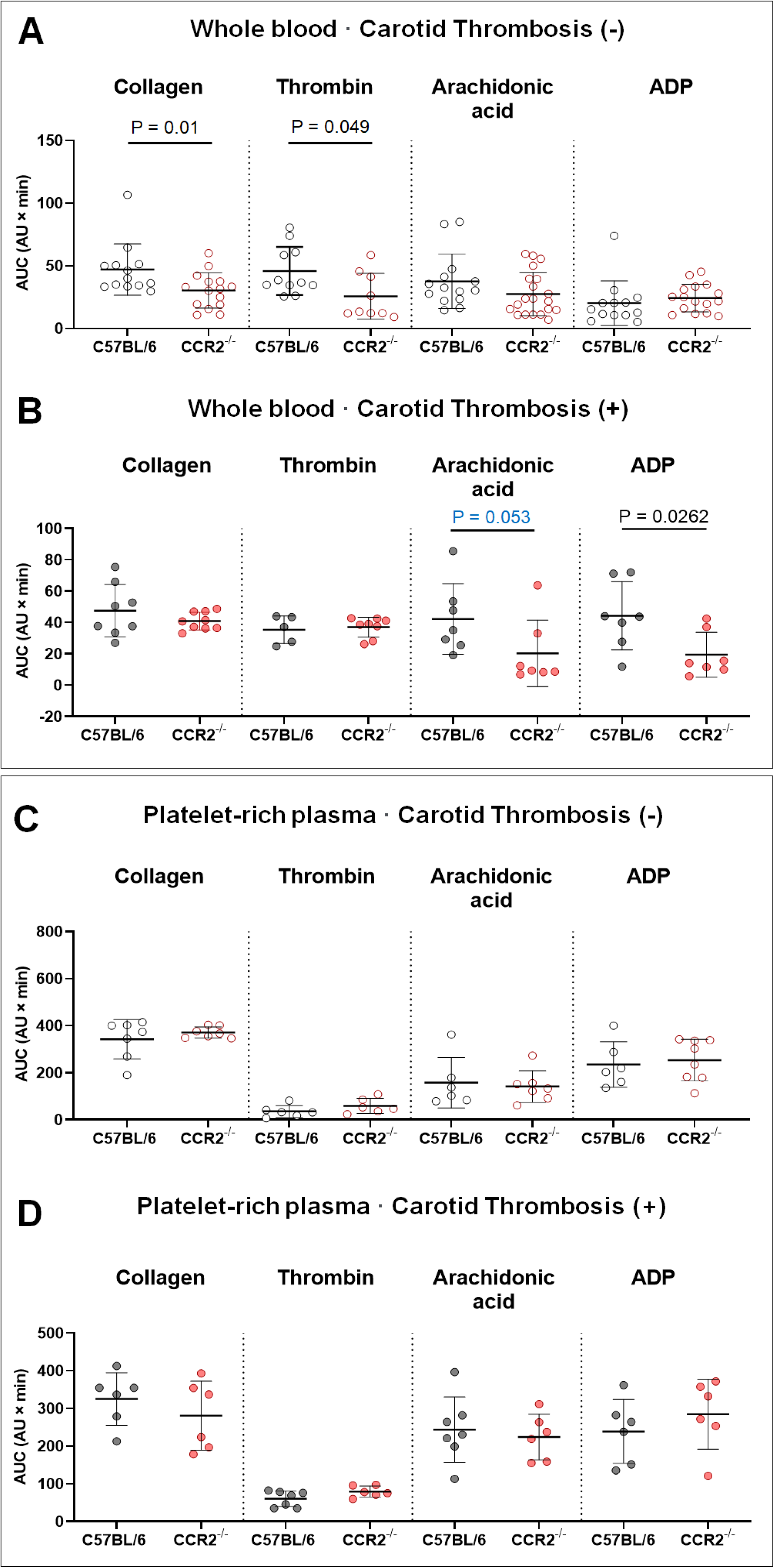
Platelet function tests using whole blood, as opposed to leukocyte-deficient platelet-rich plasma, show lower platelet aggregability in CCR2^-/-^ mice than in C57BL/6 mice. **A-D**. Area under the curve (AUC) values (mean ± SD) of collagen-, thrombin-, arachidonic acid-, or adenosine diphosphate (ADP)-induced platelet aggregation curve obtained using whole blood (**A** and **B**) or platelet-rich plasma (**C** and **D**) of mice with (**B** and **D**) or without (**A** and **C**) FeCl_3_-mediated carotid thrombosis. Graphs show mean ± SD. P values are from the Mann-Whitney U test.

Of note, platelet function tests using blood samples without monocytes, i.e. platelet-rich plasma without WBCs and RBCs, from mice with (Fig. 2C) or without (Fig. 2D) thrombosis showed no significant inter-group differences, although the response to thrombin showed slightly lower slope values in C57BL/6 mice than in CCR2^-/-^ mice.

### Circulating MPAs increased after acute carotid artery thrombosis in C57BL/6 mice but not CCR2^-/-^ mice

Immunofluorescent staining of circulating platelets and leukocytes (Fig. 3A) showed that both MPAs and NPAs were significantly increased in the blood of C57BL/6 mice at 30 minutes after carotid thrombosis, compared with non-thrombosed mice. In thrombosed CCR2^-/-^ mice, however, only NPAs increased after thrombosis, and the numbers were higher when compared with thrombosed C57BL/6 mice. These microscopy imaging-based quantifications were corroborated by flow cytometry analyses (Fig. 3B). Monocytes in the MPAs of C57BL/6 mice, particularly in thrombosed mice, were predominantly (1.277±0.5%) Ly6C-high, whereas free monocytes unbound to platelets were mostly Ly6C-low. In additional experiments, adding collagen, AA, or ADP (Supplementary Fig. 3) to blood samples elevated MPAs and NPAs in non-thrombosed C57BL/6 mice but only NPAs in non-thrombosed CCR2^-/-^ mice, as quantified by using fluorescent microscopy images. These leukocyte–platelet aggregates were further increased after thrombosis in both strains. Post-thrombosis NPA numbers in the agonist-treated blood samples did not differ significantly between C57BL/6 and CCR2^-/-^ mice, suggesting a catch-up rise occurred in C57BL/6 mice.

**Fig 3.**
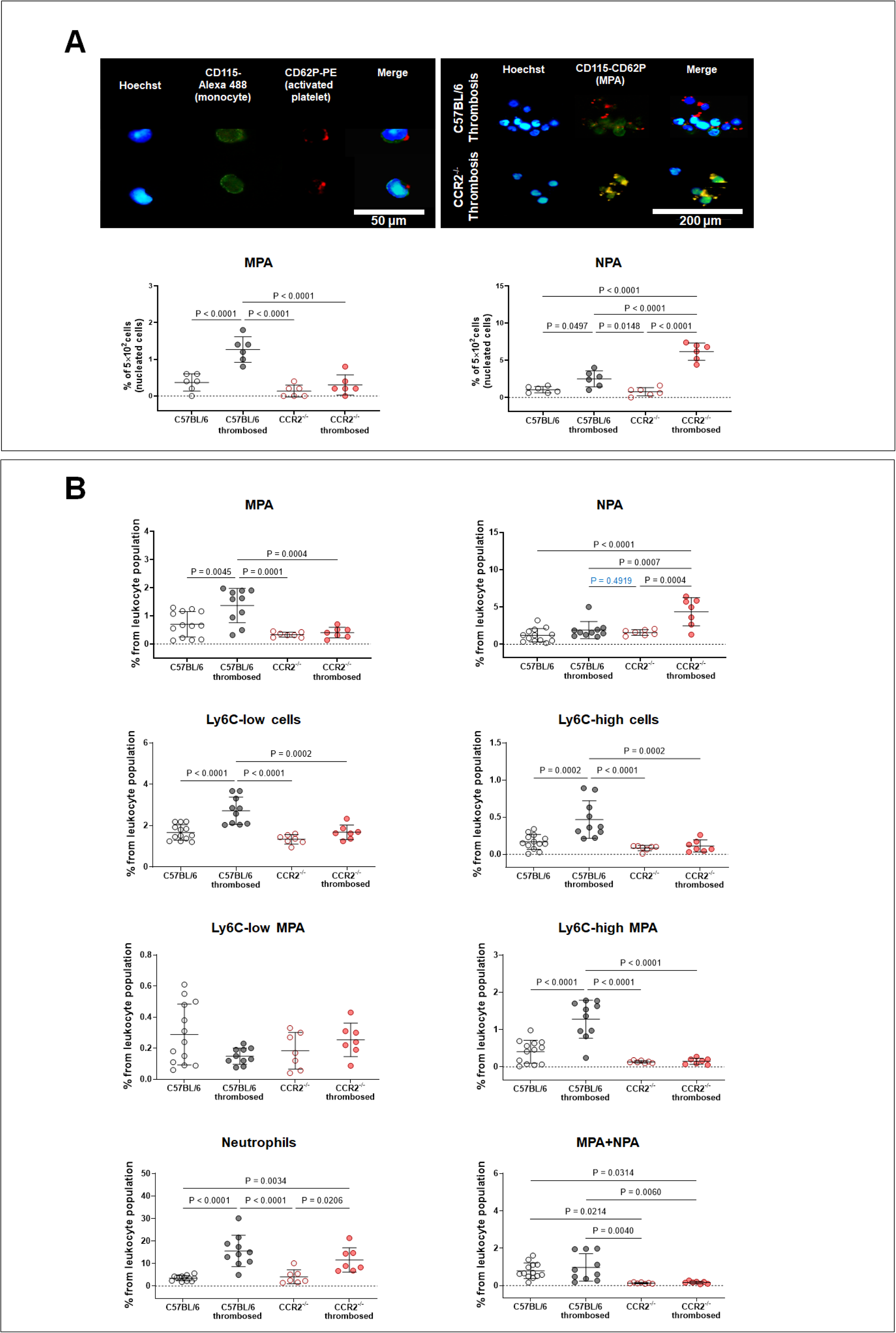
Circulating monocyte–platelet aggregates (MPA) increase after carotid artery thrombosis in C57BL/6 mice but not in CCR2^-/-^ mice. **A-B**. Microscopy imaging (**A**) and flow cytometry (**B**) quantification of (Ly6G- and CD62-positive) MPA and (DAPI- and CD62-positive) neutrophil–platelet aggregates (NPA) in whole blood obtained 30 minutes after FeCl_3_-mediated carotid thrombosis. MPA levels are higher in C57BL/6 mice than in CCR2^-/-^ mice. Also note that the post-thrombosis increase in NPA levels is more prominent in CCR2^-/-^ mice than in C57BL/6 mice, in the absence of the platelet aggregating agents. Scale bars = 50 and 200 μm. Graphs show mean ± SD. P values are from the One-way ANOVA and Tukey’s *post hoc* tests.

### Blood levels of FXIII and monocyte levels of FXIII-A rose after acute carotid artery thrombosis in C57BL/6 mice but not CCR2^-/-^ mice

Coagulation assays in plasma (Fig. 4) showed that PT and aPTT did not differ significantly between C57BL/6 mice and CCR2^-/-^ mice regardless of arterial thrombosis, nor did fibrinogen levels or bleeding times (Supplementary Fig 4). In the intrinsic pathway, there were no significant inter-group differences in FXII levels. However, FIX levels were higher in thrombosed C57BL/6 mice than in non-thrombosed C57BL/6 mice. This thrombosis-related increase in FIX levels was also observed in CCR2^-/-^ mice. Thus, FXI levels at 30 minutes after thrombosis did not differ significantly between C57BL/6 mice and CCR2^-/-^ mice. In the extrinsic pathway, tissue factor levels did not significantly differ across groups. Yet, FVII levels were significantly higher in thrombosed C57BL/6 mice compared to non-thrombosed C57BL/6 mice; CCR2^-/-^ mice also exhibited this thrombosis-related rise in FVII levels. Tissue factor levels therefore did not differ significantly between C57BL/6 mice and CCR2^-/-^ mice 30 minutes after. In the common pathway, FX and TAT levels showed no significant inter-group differences. Interestingly, FXIII levels were higher in thrombosed C57BL/6 mice than in non-thrombosed C57BL/6 mice, but this finding did not appear in CCR2^-/-^ mice. In sum, FXIII levels were significantly, albeit slightly, higher in C57BL/6 mice than in CCR2^-/-^ mice 30 minutes after thrombosis. Neither FXIIIa activity, FXa activity, nor thrombin activity showed significant inter-group differences.

**Fig 4.**
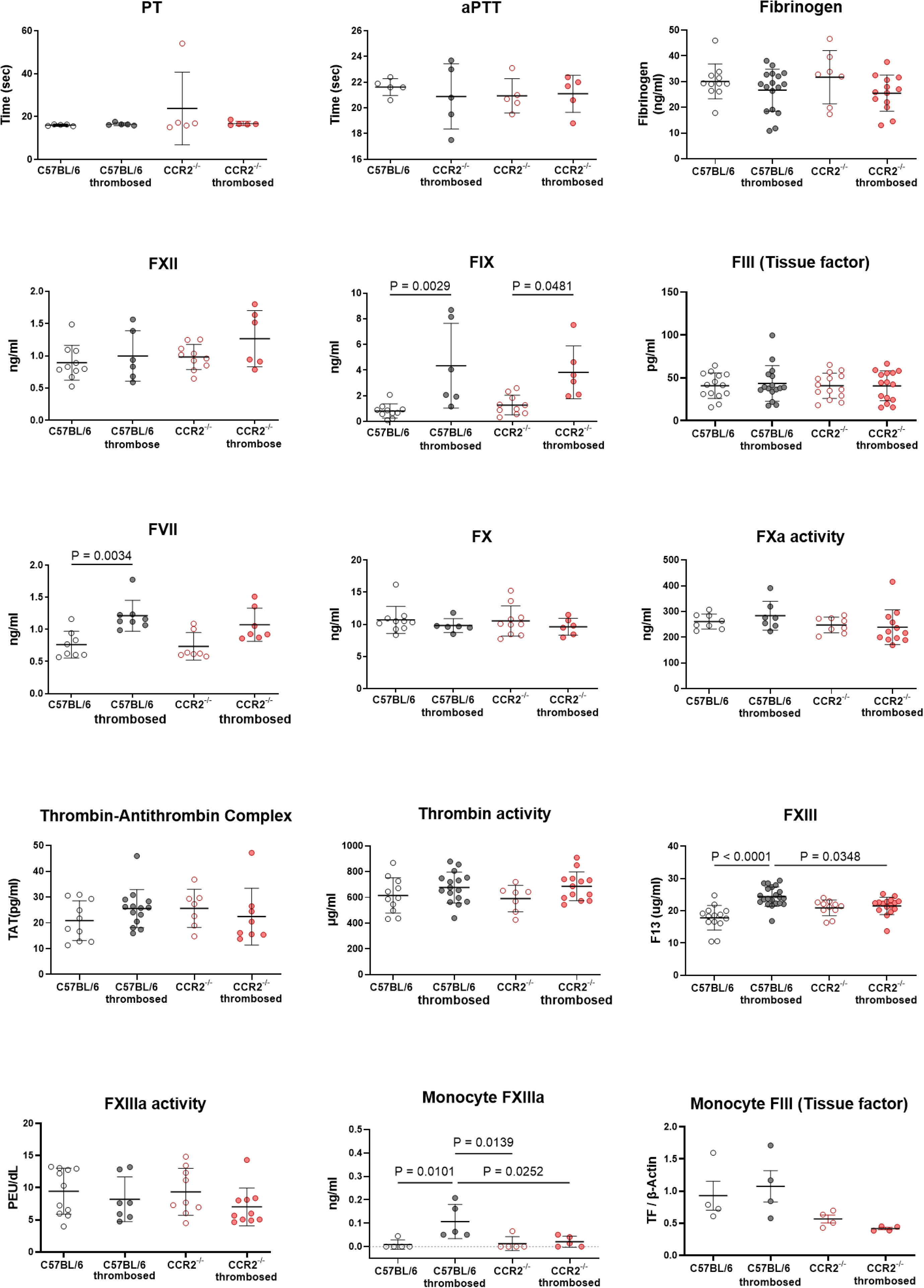
Coagulation assays show that blood levels of FXIII and monocyte levels of FXIII-A rise after carotid artery thrombosis in C57BL/6 mice but not in CCR2^-/-^ mice. Graphs show mean ± SD. P values are from the One-way ANOVA and Tukey’s *post hoc* tests.

Additionally, coagulation assays using isolated monocytes (Fig. 4) showed that FXIII-A levels were distinctly higher in thrombosed C57BL/6 mice than in non-thrombosed ones. This was not the case for tissue factor.

### MicroCT imaging and histology showed that carotid artery thrombi in CCR2^-/-^ mice were smaller, more porous, and had less fibrin cross-linking than those in C57BL/6 mice

To corroborate *in vitro* study results, we performed microCT-based thrombus imaging at 30 minutes after carotid thrombosis in 49 mice. Thrombus volume was significantly smaller in CCR2^-/-^ mice (0.112±0.002 mm^3^, n=22) than in C57BL/6 mice (0.125±0.007 mm^3^, n=27; *P* < 0.0001) (Fig. 5A and B). Histology (Fig. 5C and D) corroborated the imaging findings regarding the inter-group difference in thrombus volume. Moreover, CCR2^-/-^ mice, compared to C57BL/6 mice, had more porous thrombi (Fig. 5C) with relatively sparse fibrin cross-linking (Fig. 5D).

**Fig 5.**
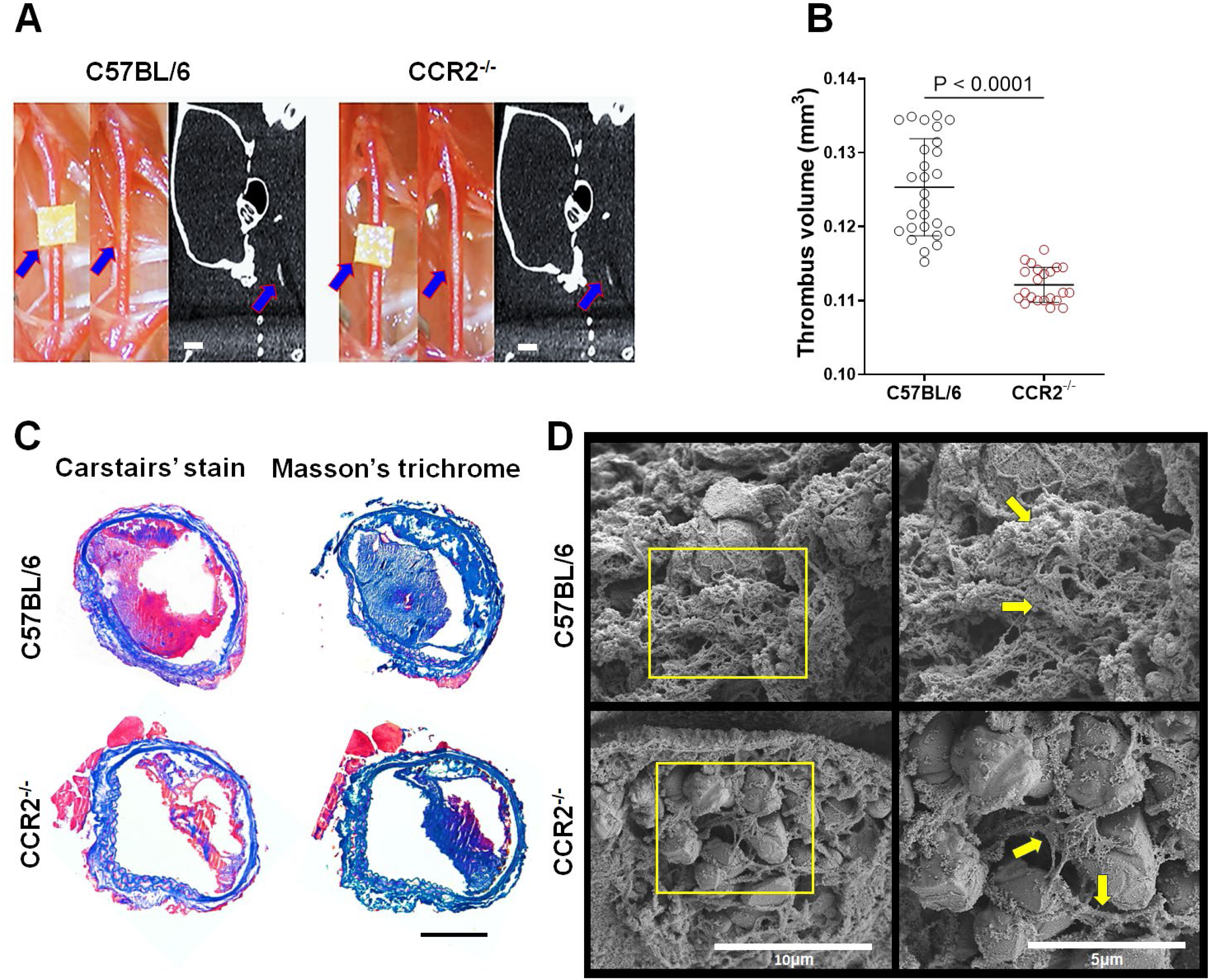
CCR2^-/-^ mice have smaller, more porous thrombi with less fibrin-crosslinking, compared to C57BL/6 mice. **A-B**. Representative microCT images (**A**) and grouped quantification data for all animals (**B**), showing significantly lower thrombus volume in CCR2^-/-^ mice than in C57BL/6 mice at 30 minutes after FeCl_3_-mediated carotid thrombosis (arrows). Scale bars = 1 mm. **C-D**. Carstair’s staining and Masson’s trichrome staining showing smaller, more porous thrombi (**C**; Scale bars = 100 μm), with relatively sparse fibrin-crosslinking (arrows in **D**, scanning electron microscopy) in CCR2^-/-^ mice, compared to C57BL/6 mice. Graphs show mean ± SD. P values are from the Mann-Whitney U test.

### tPA-mediated thrombolysis was faster in CCR2^-/-^ mice and CCR2-siRNA-treated mice, compared to control C57BL/6 mice

In thrombolysis experiments, serial microCT thrombus imaging was performed at baseline (i.e., 30 minutes after carotid thrombosis) and 1, 2, 3, and 24 hours after intravenous tPA treatment that was initiated after baseline imaging (Fig. 6). Post-tPA thrombus volume at each time-point, except for at 24 hours, was significantly lower in CCR2^-/-^ mice than in C57BL/6 mice (Fig. 6A), with or without adjusting for the baseline volume difference. Thrombus volumes at 1 and 2 hours after tPA therapy in CCR2^-/-^ mice were respectively similar to those at 2 and 3 hours in C57BL/6 mice.

**Fig 6.**
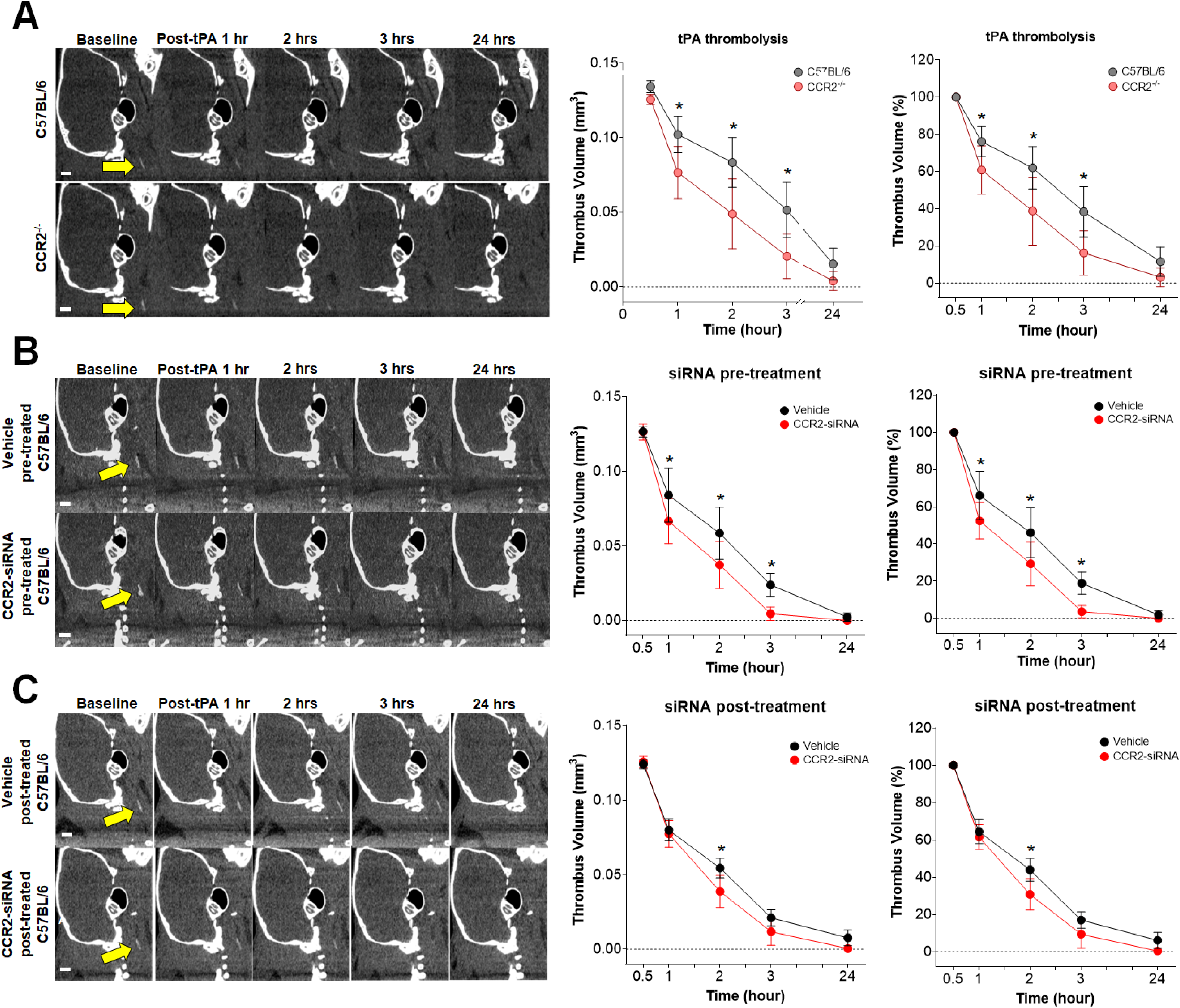
CCR2^-/-^ or CCR2 siRNA facilitates tissue plasminogen activator (tPA)-mediated thrombolysis. **A-C**. Representative serial microCT images showing tPA-mediated thrombus volume reduction (left panels) and grouped quantification data for all animals without (left graphs) or with (right graphs) adjustment for pre-tPA (baseline) thrombus volumes (mean ± SD). Note the effects of CCR2^-/-^ (**A**), CCR2-siRNA pretreatment (**B**), and CCR2-siRNA post-treatment (**C**) on tPA therapy, which was initiated about 30 minutes after FeCl_3_-mediated carotid artery thrombosis (arrows). *P<0.05 for the inter-group difference within each time-point: linear mixed models with random intercepts and pairwise *post hoc* tests with Sidak’s adjustment for multiple comparisons. Scale bars = 1 mm.

In two additional sets of thrombolysis experiments using C57BL/6 mice only (Fig. 6B and C), serial microCT imaging was performed, as described above, at baseline and serially after tPA therapy, but with either daily CCR2-siRNA (vs. vehicle) pretreatment for 3 days (Fig. 6B) or a single dose of CCR2-siRNA (vs. vehicle) post-treatment (Fig. 6C). tPA-mediated reduction of thrombus volume was largest in C57BL/6 mice that were pretreated with CCR2-siRNA, as these showed smaller thrombus volumes at 1, 2, and 3 hours, compared to the vehicle group. In the CCR2-siRNA post-treatment group, thrombus volume was decreased at 2 hours only, compared to the vehicle group.

The last set of thrombolysis experiments (Fig. 7) showed lower baseline thrombus volumes in C57BL/6 mice pretreated with clopidogrel (0.092±0.005 mm^3^) and CCR2^-/-^ mice pretreated with clopidogrel (0.085±0.007 mm^3^) than in C57BL/6 mice (0.132±0.007 mm^3^) and CCR2^-/-^ mice (0.117±0.003 mm^3^). tPA-mediated reduction of thrombus volume was larger in the two CCR2^-/-^ mouse groups (pretreated with either clopidogrel or saline), smaller in the clopidogrel-pretreated C57BL/6 mouse group, and smallest in the saline-pretreated C57BL/6 mouse group. Taken together, these findings indicate that clopidogrel had a greater effect on inducing thrombus formation with smaller sizes after FeCl_3_ application, while CCR2^-/-^ had a greater effect on dissolving thrombus faster after tPA administration.

**Fig 7.**
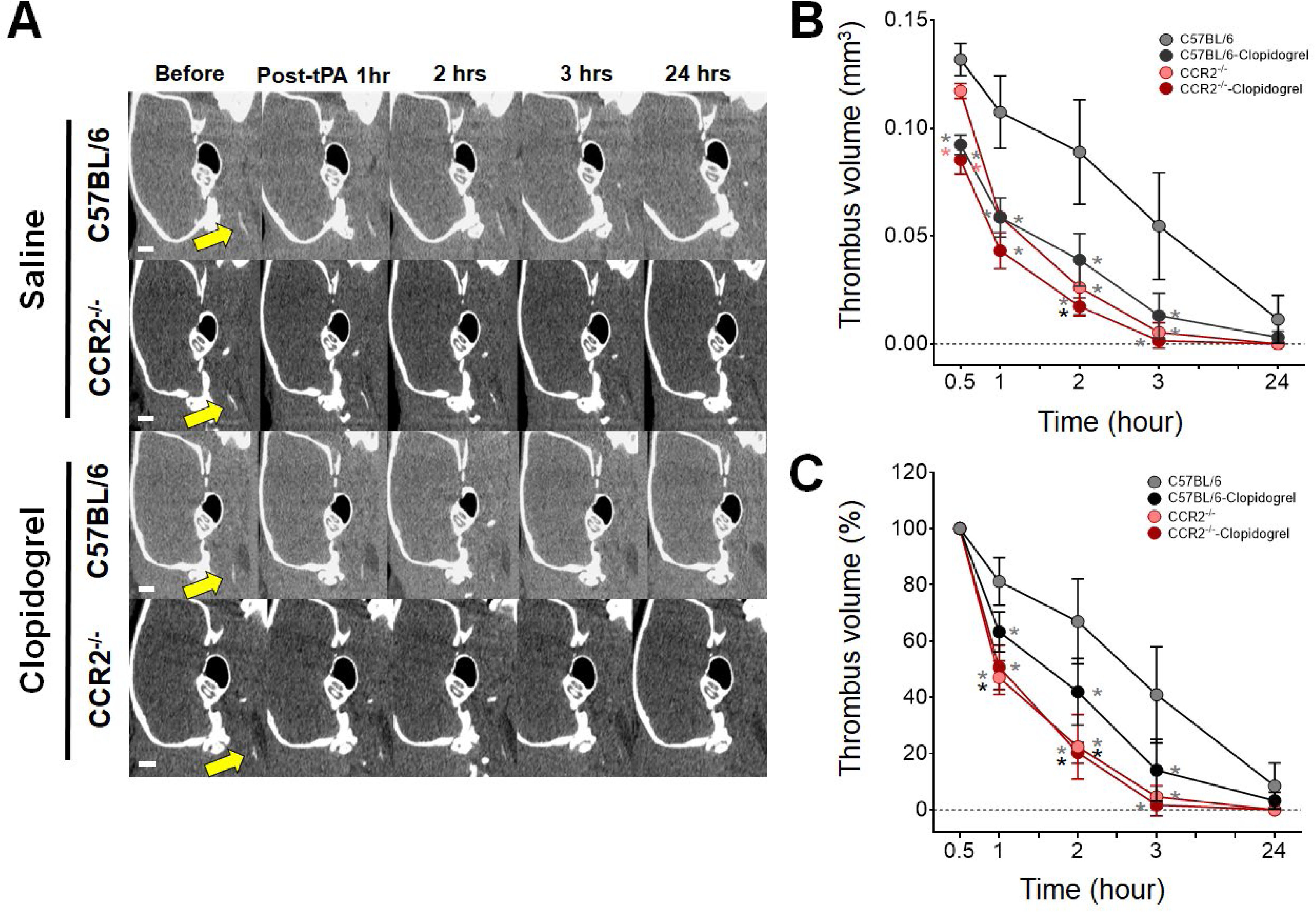
CCR2^-/-^, compared with clopidogrel pretreatment, shows less reduction in initial thrombus volume but greater enhancement in tissue plasminogen activator (tPA)-mediated thrombolysis. **A-C**. Representative serial microCT images showing tPA-mediated reduction of thrombus volume (**A**) and grouped quantification data for all animals without (**B**) or with (**C**) adjustment for pre-tPA (baseline) thrombus volumes (mean ± SD). Note how CCR2^-/-^ and/or 7-day clopidogrel treatment affect baseline thrombus volumes and tPA-mediated thrombus volume reduction. tPA therapy was initiated ∼1 hour after FeCl_3_-mediated carotid artery thrombosis (arrows). Gray, black, and orange * indicate P < 0.05 vs. the C57BL/6, C57BL/6-clopidogrel, and CCR2^-/-^ mouse groups, respectively, at each time-point: linear mixed models with random intercepts and pairwise *post hoc* tests with Sidak’s adjustment for multiple comparisons. Scale bars = 1 mm.

## Discussion

This study demonstrates that CCR2^-/-^-mediated monocyte deficiency reduces platelet aggregation, probably in part by decreasing MPAs (which have known prothrombotic effects), thereby a) generating smaller, more porous thrombi after FeCl_3_ application on the carotid artery and thus b) facilitating intravenous tPA-mediated thrombolysis *in vivo*. Moreover, CCR2 knockouts had lower post-thrombosis FXIII levels in blood and monocytes, which could explain the sparser fibrin cross-linking during arterial thrombosis and may be another important mechanism of these antithrombotic effects.

We previously developed a microCT-based direct thrombus imaging technique, using thrombus-seeking gold nanoparticles, and have shown its preclinical utility for a) quantitatively detecting arterial thrombus^11,12^ and b) serially monitoring prothrombotic^15^ and antithrombotic^23^ effects. In the present study, this *in vivo* imaging technique showed acute carotid thrombi to be significantly smaller in CCR2^-/-^ mice than in C57BL/6 mice. In addition, histology showed that CCR2^-/-^ mice had a smaller, more porous thrombus, compared with C57BL/6 mice. These findings provide, to our knowledge, the first direct *in vivo* evidence that monocytes play a prothrombotic role in arterial clots.

The antithrombotic phenotypes of CCR2^-/-^ mice (vs. C57BL/6 mice) can be attributed to relatively attenuated platelet aggregation, as evidenced by platelet function tests performed using whole blood. Tests in platelet-rich plasma showed no significant platelet aggregability differences between the two strains, a result that indicates blood cells other than platelets also contribute to the observed differences. When leukocyte–platelet aggregates—biomarkers of platelet activity and predictors of cardiovascular outcomes^25,26^—were quantified using immunofluorescent staining and flow cytometry, both MPA and NPA numbers were elevated in the blood of C57BL/6 mice after carotid thrombosis; however, in CCR2^-/-^ mice, only NPA numbers were increased.

Intravital microscopy and flow cytometry demonstrated that circulating monocytes and neutrophils accumulated in carotid thrombus. These leukocytes could have been passively entrapped^27^ due to flow disturbance near the clot. Yet, notably high thrombus-blood ratios of monocytes and neutrophils (vs. those of lymphocytes) suggest these leukocytes were actively recruited. Gene expression analysis also showed upregulation of cytokines (CCL2, CCL5, CCL7, CXCL2, CX3CR1, IL-6, and IL-1b), which may have facilitated leukocyte recruitment into thrombus^28–30^ in the lesional (vs. non-thrombosed contralateral) carotid arteries. Thus, less monocyte recruitment to arterial thrombi as well as lower monocyte counts in the blood may have decreased monocyte–platelet interaction and platelet aggregability during acute arterial thrombosis in CCR2^-/-^ mice, compared to C57BL/6 mice.

Early during thrombogenesis, FXIIIa cross-links fibrin monomers and hence stabilizes the fibrin clot.^31–33^ Such cross-linking causes fiber compaction, which shrinks pores within individual fibers and strengthens the rigidity of the condensed clot,^34^ thereby limiting tPA diffusion through the fibers.^35^ Another mechanism of FXIIIa-mediated thrombus protection from fibrinolysis involves incorporating the fibrinolysis inhibitor alpha 2-antiplasmin into the clot.^36^ We found that, compared with C57BL/6 mice, CCR2^-/-^ mice had lower post-thrombosis levels of monocyte-derived FXIII-A, which is the cellular form of FXIII synthesized and non-proteolytically activated by increases in intracellular Ca^2+^ concentrations.^37^ This result, in conjunction with the intravital microscopy and flow cytometry studies showing that monocytes were actively incorporated into the fibrin mesh of model thrombi under shear stress, suggests that decreased fibrin cross-linking in thrombi may contribute to the faster tPA-mediated fibrinolysis observed in CCR2^-/-^ mice (vs. C57BL/6 mice). A recent *in vitro* study^37^ showed that cytokine-mediated stimulation of monocytes i) exposed FXIII-A on the cell surface and ii) reduced tPA-mediated degradation of FXIII-deficient thrombi ∼two-fold. Taken together, these data indicate that monocytes act as a vehicle to deliver FXIII-A into arterial thrombi, thereby stabilizing the thrombus against fibrinolytic degradation.^37^

Pretreating C57BL/6 mice with CCR2-siRNA (vs. saline) for 3 days, as was the case for the CCR2^-/-^ vs. wild-type genotype, led to more efficient thrombolysis by tPA administered within the clinically approved therapeutic time window (∼4.5 hours). These results suggest the translational potential of modulating monocytes to facilitate thrombolysis in patients, although post-treatment with single-dose CCR2-siRNA at 30 minutes after thrombosis was less effective than the pretreatment. Compared with saline-pretreated C57BL/6 mice, saline-pretreated CCR2^-/-^ mice had ∼10% smaller thrombi and clopidogrel-pretreated CCR2^-/-^ mice had ∼36% smaller thrombi. These findings may suggest that CCR2^-/-^ and inhibiting the platelet P2Y12 ADP receptor could have non-redundant effects on arterial thrombus formation. However, clopidogrel-pretreated C57BL/6 mice had ∼31% smaller thrombi than saline-pretreated C57BL/6 mice. Clopidogrel effects therefore appear to supersede CCR2 knockout effects on thrombus formation in our experimental setting. Nevertheless, this ceiling effect was not observed in tPA-mediated thrombolysis: the thrombolysis rate was higher in saline-pretreated CCR2^-/-^ mice and clopidogrel-treated CCR2^-/-^ mice, lower in clopidogrel-pretreated C57BL/6 mice, and lowest in saline-pretreated C57BL/6 mice.

Developing more potent antithrombotic therapies that inhibit the activity of either platelets or coagulation factors has been limited by the consequently elevated risk of hemorrhagic complications.^38,39^ Future studies should investigate whether adjusting thromboinflammation via an alternative approach, such as inhibiting CCR2, can improve both safety and efficacy in treating ischemic infarction. We showed that PT, aPTT, fibrinogen levels, and bleeding time did not differ between CCR2^-/-^ mice and C57BL/6 mice. In addition, clopidogrel reduced baseline thrombus volume more, but CCR2^-/-^ better facilitated tPA-mediated thrombolysis. Further investigation is required to study the clinical potential of combining CCR2 inhibition with conventional antithrombotic or thrombolytic therapy in ischemic stroke.

Our study has limitations. First, the molecular mechanism underlying the monocyte deficiency-related reduction in FXIII levels in blood and monocytes still needs to be elucidated. Second, it is unclear whether the increased NPA in CCR2^-/-^ mice (vs. C57BL/6 mice) was (at least partly) compensating for the MPA reduction-related dysfunctional platelet aggregability. Third, our murine experiments need to be validated by clinical research. Nevertheless, by demonstrating the antithrombotic and tPA-facilitating effects of monocyte targeting in a mouse model of carotid thrombosis, our study may provide the groundwork for future therapeutic innovations.

## Non-standard Abbreviations and Acronyms

aPTT: activated partial thromboplastin time
ADP: adenosine diphosphate
A6: amplitude at 6 minutes in Ohm
AA: arachidonic acid
AUC: area under the aggregation curve
BV: Brilliant Violet
CCL2: CC chemokine ligand 2
CCR2: chemokine receptor 2
CCA: common carotid artery
ELISA: enzyme-linked immunosorbent assay
fib-GC-AuNPs: fibrin-targeted glycol-chitosan-coated gold nanoparticles
LPA: leukocyte-platelet aggregates
MT: Masson’s trichrome
microCT: micro-computed tomography
MPA: monocyte-platelet aggregates
NPA: neutrophil-platelet aggregates
PPP: platelet-poor plasma
PRP: platelet-rich plasma
PT: prothrombin time
SEM: scanning electron microscope
SD: standard deviation
TAT: thrombin-antithrombin complex
tPA: tissue plasminogen activatorIntroduction

## Supplemental Material

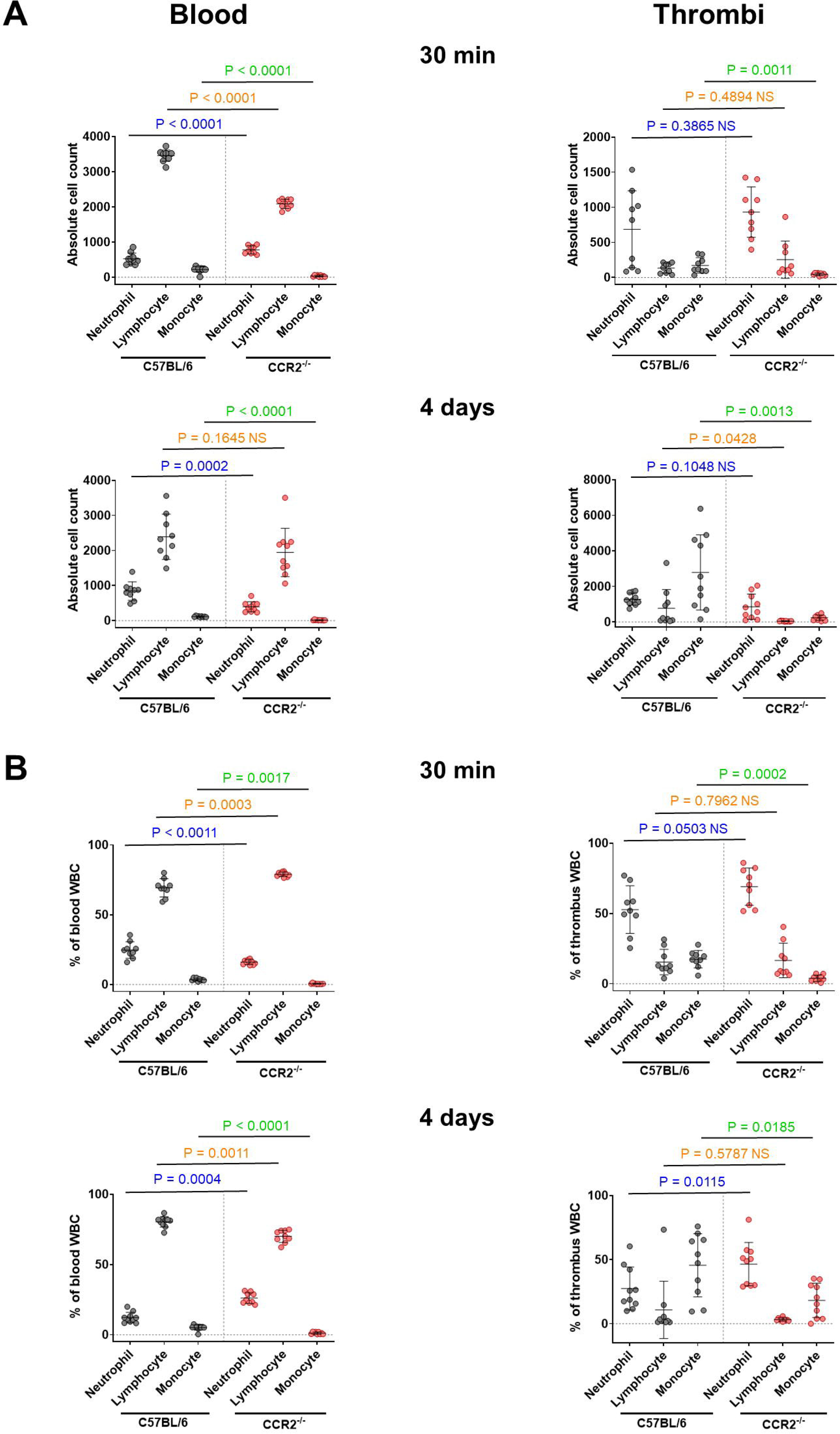

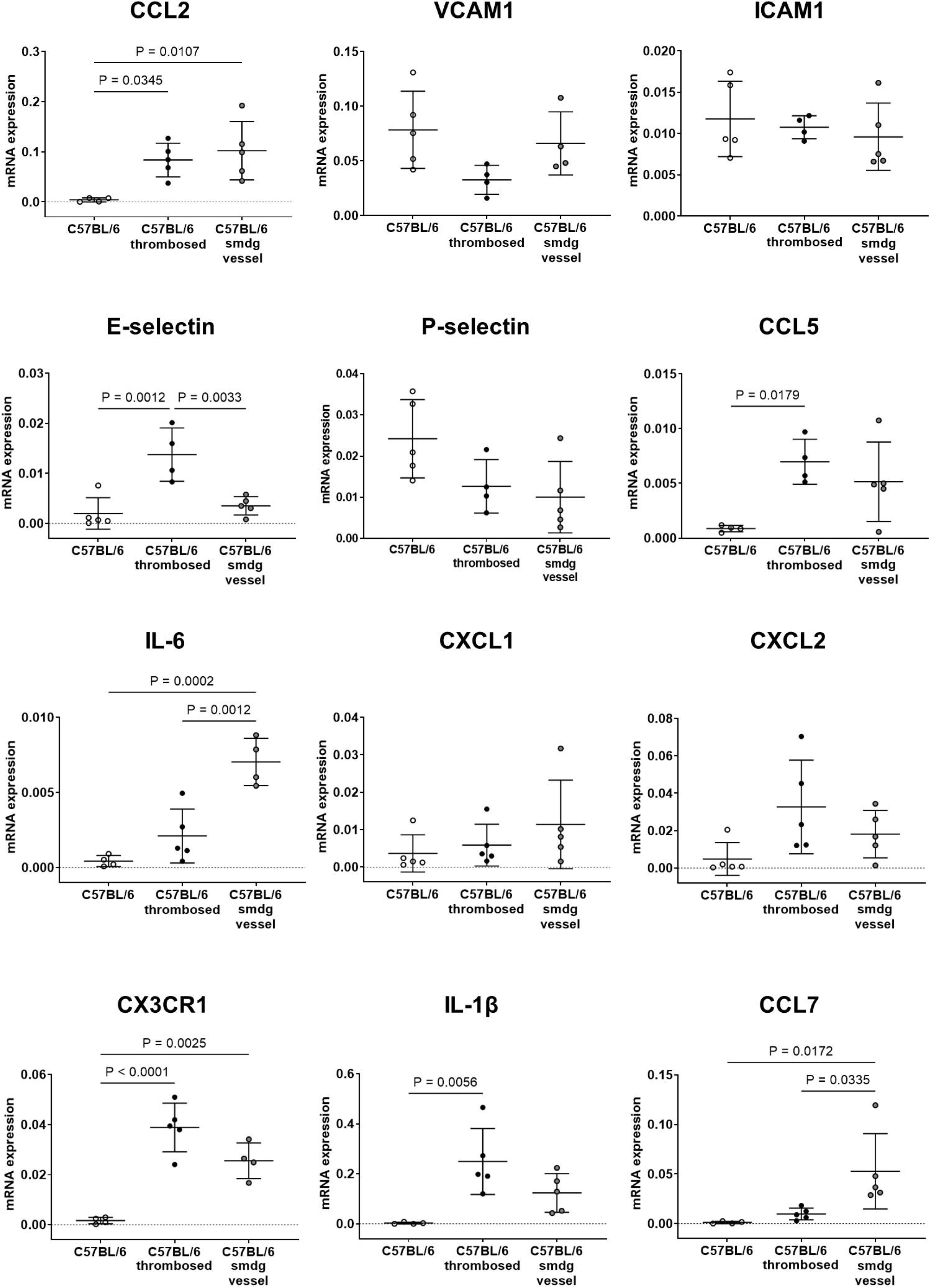

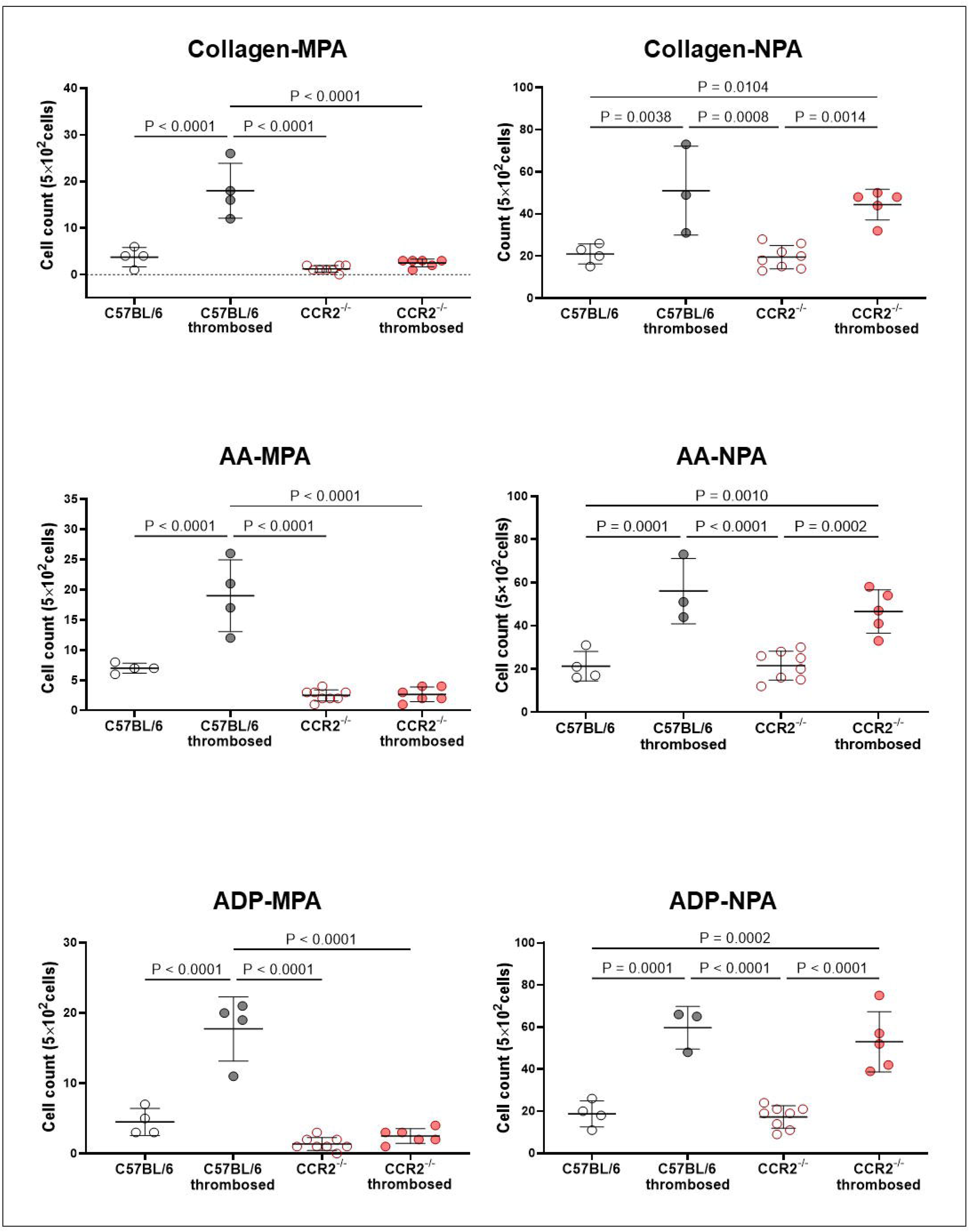

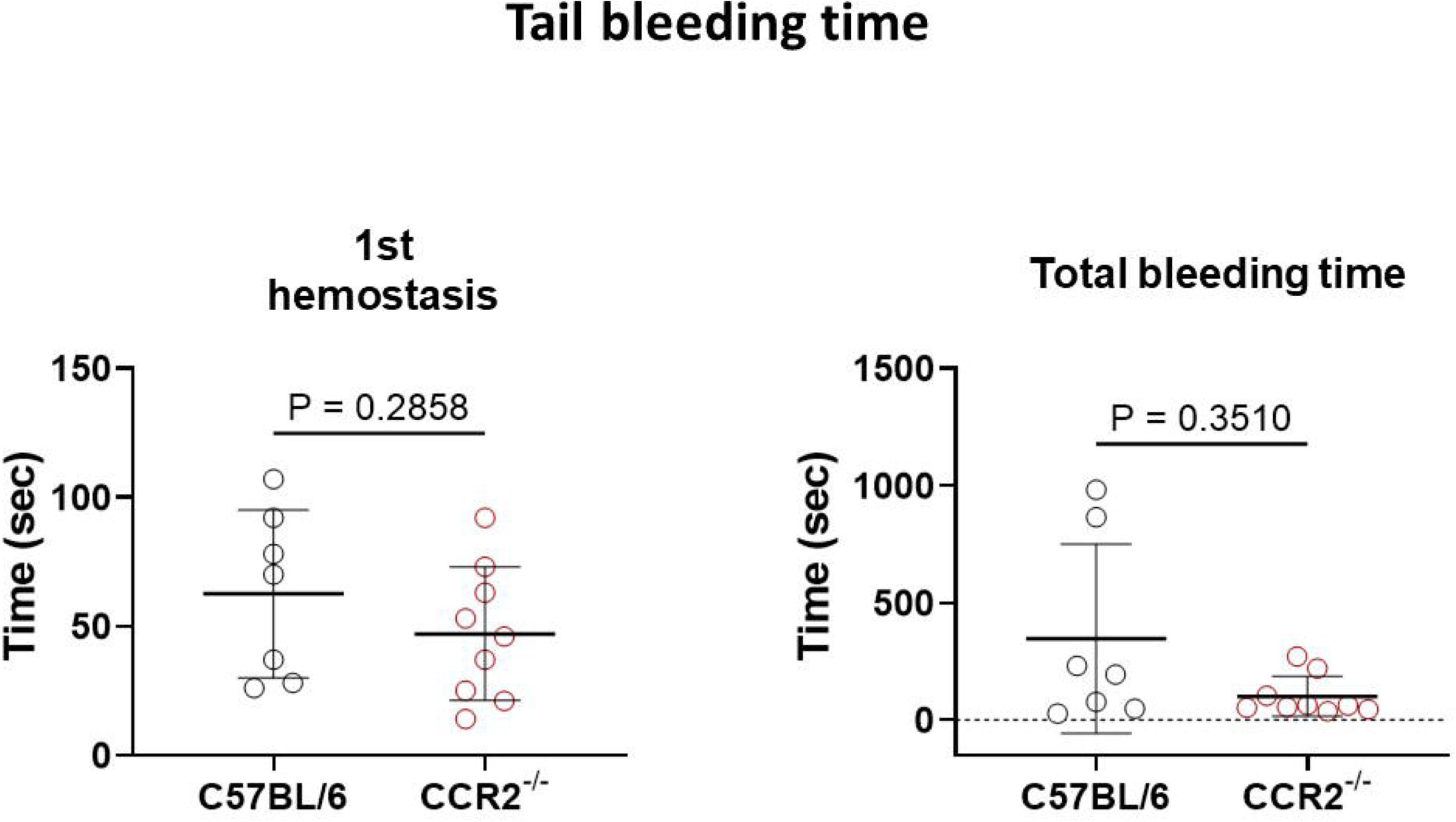

