## supplementary material for "Inhibiting monocyte migration reduces arterial thrombosis and facilitates thrombolysis"

Department of Neurology, Dongguk University Ilsan Hospital, Goyang, Republic of Korea (H.J.J., J.K., H.K., T.K., D.-E.K.). National Priority Research Center for Stroke, Goyang, Republic of Korea (H.J.J., J.K., H.K., T.K., J.C., D.-E.K.). Medical Science Research Center, Dongguk University, Goyang, Republic of Korea (J.C.). Departments of Neuroradiology and Imaging Physics, University of Texas MD Anderson Cancer Center, Houston, TX (D.S.). Center for Systems Biology, Massachusetts General Hospital and Harvard Medical School, Boston, MA, USA (S.C., M.K., M.N.). Department of Radiology, Massachusetts General Hospital and Harvard Medical School, Boston, MA, USA (M.N.). Department of Internal Medicine, University Hospital Wuerzburg, Wuerzburg, Germany (M.N.). Gordon Center for Medical Imaging, Massachusetts General Hospital, Boston, MA, USA (M.N).

^*^These authors contributed equally to this work.

**Short title: Monocytes in arterial thrombosis**

**Correspondence to:**

Dong-Eog Kim, MD, PhD

National Priority Research Center for Stroke, Dongguk University Ilsan Hospital, 27, Dongguk-ro, Ilsandong-gu, Goyang, Republic of Korea 10326,

&

Matthias Nahrendorf , MD, PhD

Center for Systems Biology, Massachusetts General Hospital, Richard B. Simches Research Center, 185 Cambridge Street, Suite 5.210, Boston, MA 02114, USA,

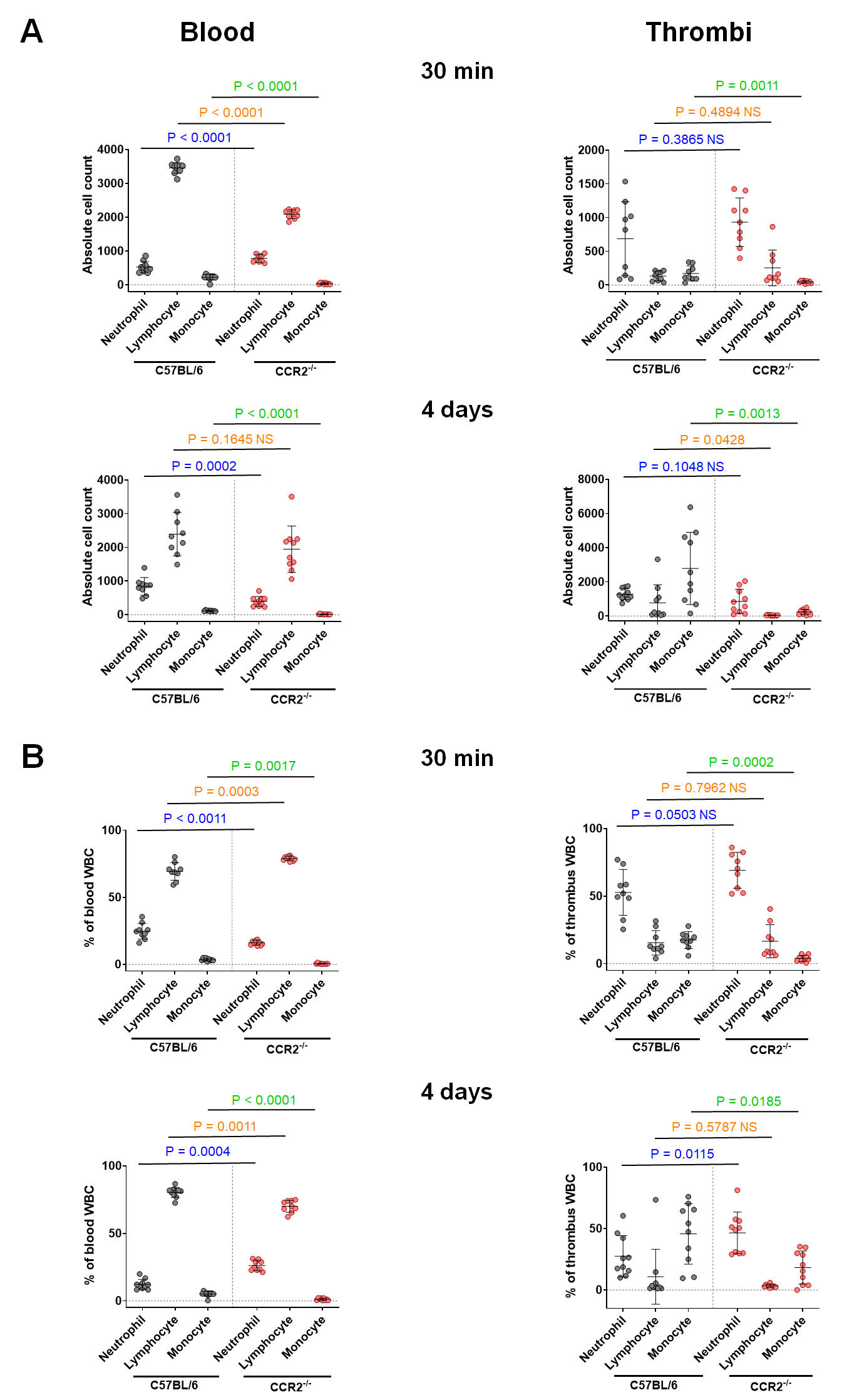

**Supplementary Figure 1.** Scatter plots for Fig. 1 data. Graphs show mean ± SD. P values are from the Mann-Whitney U test.

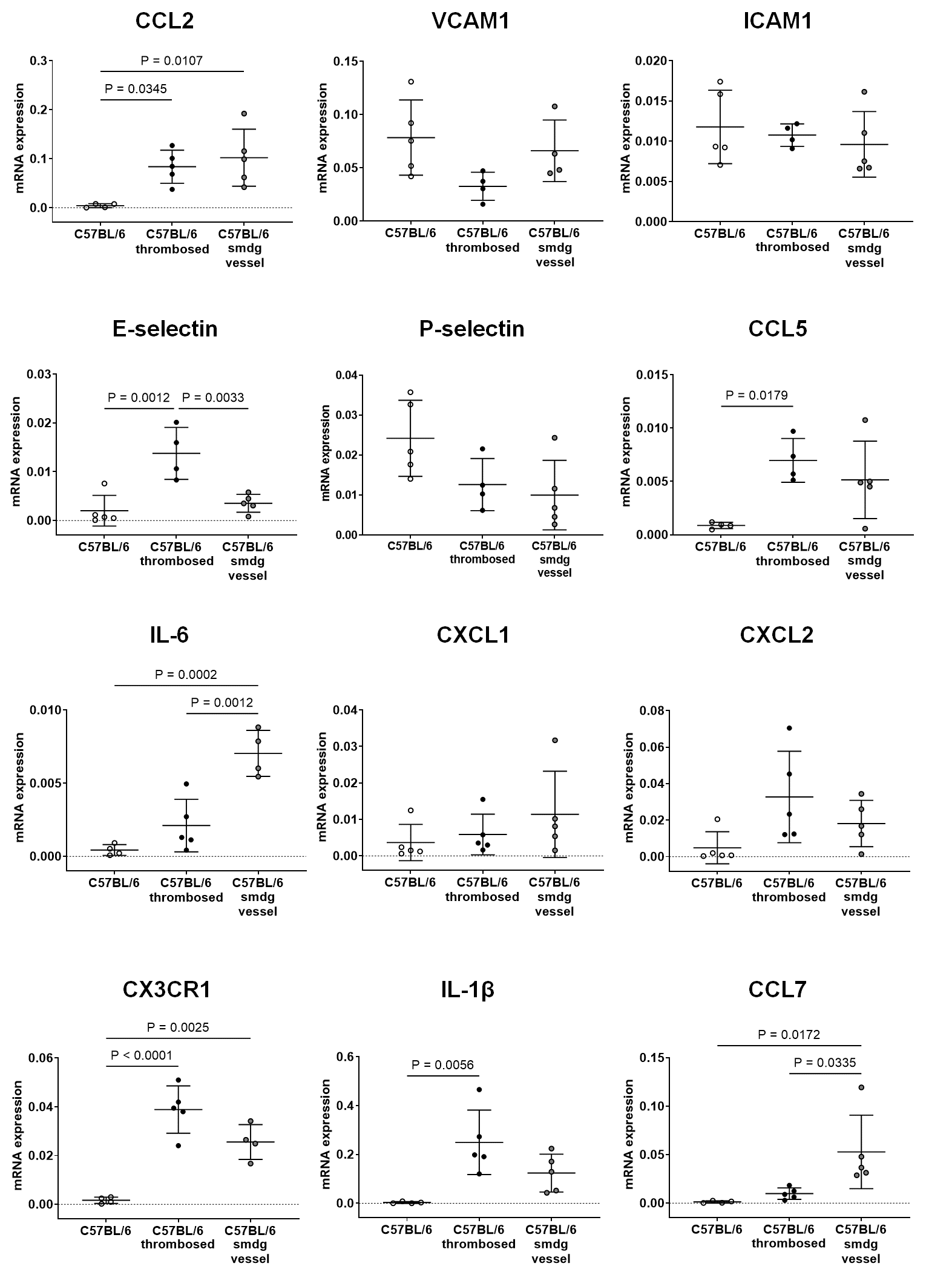

**Supplementary Figure 2. Cytokine analysis.** Graphs show mean ± SD. P values are from the One-way ANOVA and Tukey’s *post hoc* tests. CCL2, CC chemokine Ligand 2; VCAM1, vascular cell adhesion molecule 1; ICAM1, intercellular adhesion molecule 1; E-selectin, endothelial selectin; P-selectin, platelet selectin; CCL5, CC chemokine Ligand 5; IL-6, interleukin 6; CXCL1, CXC chemokine Ligand 1; CXCL2, CXC chemokine Ligand 2; CX3CR1, CX3C chemokine receptor 1; IL-1β, interleukin 1β.

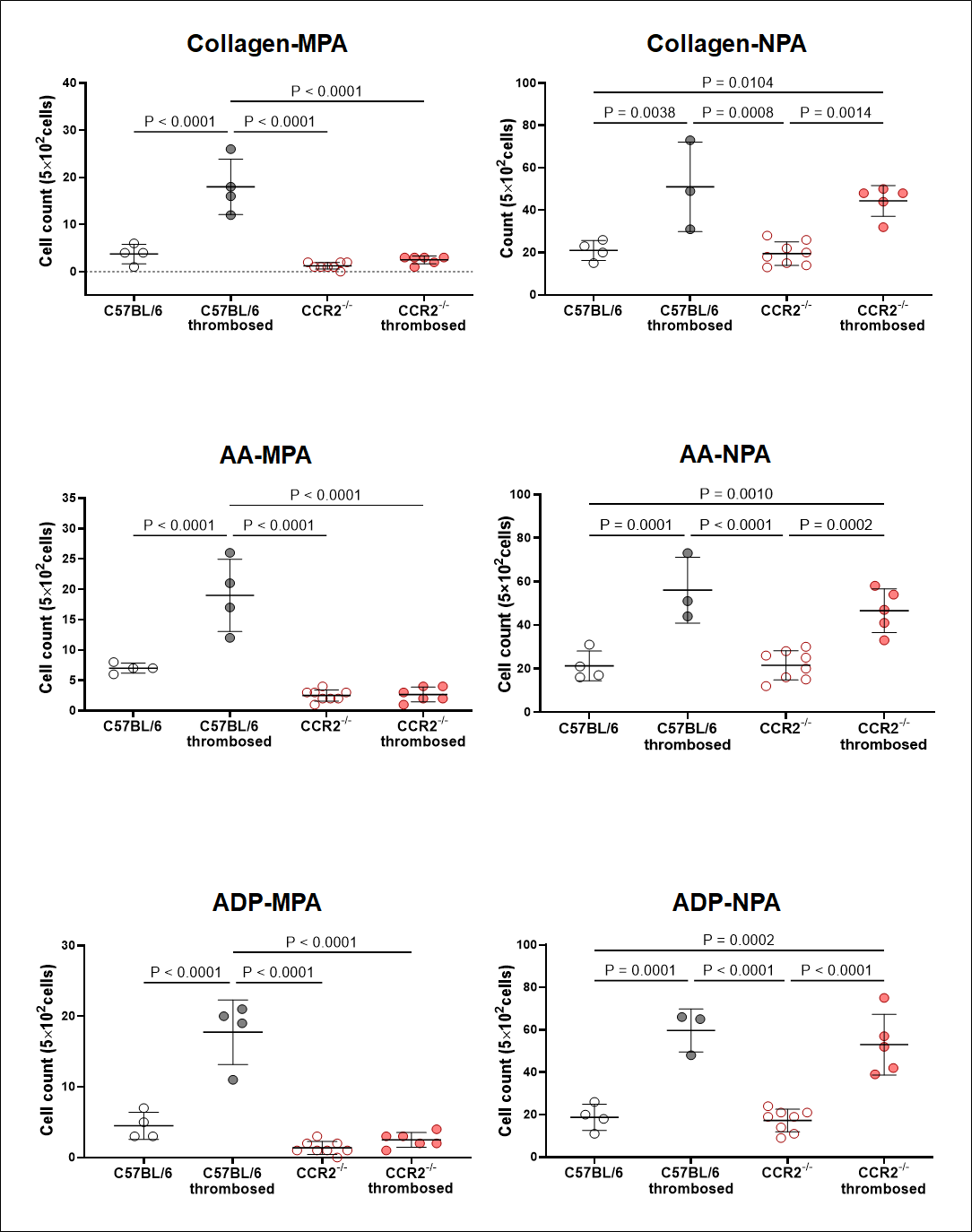

**Supplementary Fig 3.** Microscopy imaging-based quantification of monocyte–platelet aggregates (MPA) and neutrophil–platelet aggregates (NPA) after adding collagen, arachidonic acid (A), or adenosine diphosphate (ADP) to whole blood drawn from either C57BL/6 mice or CCR2^-/-^ mice with or without thrombosis. Graphs show mean ± SD. P values are from the One-way ANOVA and Tukey’s *post hoc* tests.

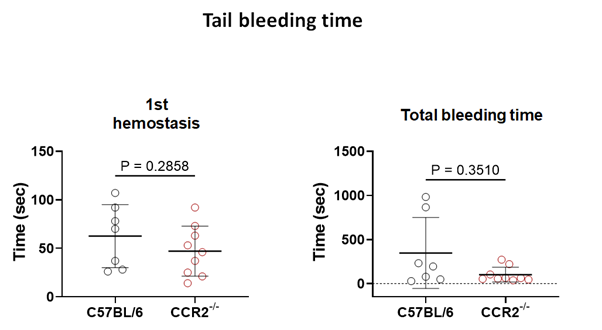

**Supplementary Fig 4. Tail bleeding time.** Graphs show mean ± SD. P values are from the Mann-Whitney U test.
